# Claudin-4, a core component of the tight-junctional complex along the collecting system, is induced in nephrotic syndrome

**DOI:** 10.1101/2022.06.01.494142

**Authors:** Valérie Olivier, Ali Sassi, Gregoire Arnoux, Regine Chambrey, Isabelle Roth, Alexandra Chassot, Khalil Udwan, Eva Dizin, Joseph M. Rutkowski, Lydie Cheval, Gilles Crambert, Carsten A. Wagner, Alain Doucet, Dominique Eladari, Solange Moll, Eric Feraille, Suresh K Ramakrishnan

**Author notes:** Corresponding authors: Eric Feraille and Suresh K Ramakrishnan Department of Cellular Physiology and Metabolism Faculty of Medicine University of Geneva CMU, 1 Rue Michel Servet CH-1211 Geneva 4 Switzerland. These authors contributed equally to this work (Valérie Olivier, Ali Sassi, Gregoire Arnoux). These authors share senior authorship (Eric Feraille, Suresh K Ramakrishnan).

## Abstract

**Background:** Nephrotic syndrome (NS) is characterized by massive sodium chloride retention. Along the kidney tubule, sodium and chloride reabsorption are coupled *via* a combination of transcellular and paracellular transport pathways. The mechanism of sodium retention in NS has been extensively studied, but the associated chloride transport pathway has not been elucidated.

**Methods:** To investigate the pathway of chloride retention in NS, we assessed the expression levels of both paracellular and transcellular components of chloride transport in the CD of POD-ATTAC mice and PAN rats, two rodent models of NS. We also used cultured mouse cortical collecting duct cells to see how overexpression or silencing of claudin-4 affect paracellular permeability. Finally, human renal biopsies were used to confirm our *in vivo* results.

**Results:** In control animals, claudin-4 was expressed at low levels in collecting duct (CD). In POD-ATTAC mice and PAN rats, claudin-4 expression was strongly increased in CD beta-intercalated cells (B-IC) and to a lesser extent in CD principal cells and was also induced in connecting tubules. Similarly, we found that claudin-4 was expressed at low levels in normal human kidneys and was dramatically increased in CD cells of nephrotic human kidneys (focal and segmental glomerulosclerosis). In parallel, the expression of pendrin, which exchanges chloride for bicarbonates in B-IC, was decreased in nephrotic compared to control animals. However, the increase in claudin-4 expression observed in NS is likely independent of pendrin abundance. Increased claudin-4 abundance is coupled with increased ENaC-dependent sodium transport. Overexpression or silencing of claudin-4 in mCCD_cl1_ cells confirmed the preferential permeability of claudin-4 to chloride over sodium.

**Conclusions:** These results suggest that during NS, transcellular Cl-/HCO ^-^ transport decreases while paracellular chloride transport *via* claudin-4 may increase along the collecting system. Paracellular chloride permeability may constitute a chloride shunt that favors Na^+^ reabsorption and opposes K^+^ secretion along the CD in NS.

**Significance Statement:** Nephrotic syndrome is a common disease characterized by massive proteinuria, hypoalbuminemia and edema due to renal sodium-chloride retention. We demonstrate for the first time an induction of claudin-4 expression indicating a partial shift from transcellular to paracellular chloride transport in the renal collecting system of nephrotic rodents. We confirmed the increased expression of claudin-4 in kidney biopsies of nephrotic patients, highlighting the translational significance of these results. Whether the paracellular pathway may represent a novel target to treat edema in nephrotic syndrome remains to be elucidated.

## Introduction

Nephrotic syndrome (NS) is characterized by massive proteinuria, hypoalbuminemia and edema secondary to renal sodium chloride retention ^1, 2^. Along the kidney tubule, sodium and chloride reabsorption are coupled *via* a combination of transcellular and paracellular transport pathways. Previous studies have shown that transcellular sodium reabsorption *via* the Epithelial Na^+^ Channel (ENaC) is enhanced along the aldosterone sensitive distal nephron (ASDN) in NS ^3, 4^. ENaC allows the apical entry of sodium by free diffusion following its electrochemical gradient in principal cells ^5^. Sodium is then released at the basolateral side of tubular cells by the Na-K-ATPase to maintain the sodium concentration gradient ^6^. The extent of sodium reabsorption relies mostly on both sodium delivery to the ASDN and amounts of active ENaC and Na,K-ATPase of tubular cells. The activation of ENaC is regulated by aldosterone, that stimulates the synthesis of the rate-limiting α-ENaC subunit, thereby increasing ENaC subunit assembly and driving the apical translocation of assembled ENaC ^7^. Aldosterone additionally induces ENaC activation *via* stimulation of the proteolytic cleavage of its γ-subunit ^4^. Conversely, other experiments suggest that γ-ENaC may be directly cleaved by protease(s) from nephrotic urine independently of aldosterone ^8^.

Under physiological conditions, chloride reabsorption by the collecting duct (CD) predominantly occurs through the transcellular pathway *via* the apical Cl^-^ / HCO_3_ ^-^ exchanger pendrin expressed in β-intercalated cells (B-IC) ^9^. A fraction of chloride reabsorption may also occur through the paracellular pathway. At least two claudin proteins, claudin-4 and claudin-8 may contribute to this paracellular chloride reabsorption along the ASDN. This hypothesis is supported by results obtained in claudin-4 or claudin-8 knockout mice, which displayed renal NaCl wasting, hypotension, hypochloremia and metabolic acidosis ^10, 11^. Paracellular chloride reabsorption may decrease the lumen negative transepithelial potential and thereby may favor transcellular sodium reabsorption.

The pathway accounting for the increased chloride reabsorption associated with sodium retention in NS has not been characterized. For this purpose, we assessed the expression levels of both paracellular and transcellular components of chloride transport in the CD of POD-ATTAC mice and PAN rats, two rodent models of NS. We found that claudin-4 abundance increased while pendrin abundance decreased in both models of NS. This increase in claudin- 4 abundance along the CD was also found in biopsies from human nephrotic kidneys. Finally, we confirmed that claudin-4 is relatively more permeable to chloride over sodium in cultured CD principal cells. These results suggest that in NS, chloride reabsorption shifts from a predominantly transcellular pathway to a paracellular chloride pathway generated by induction of claudin-4 expression. The generated paracellular chloride shunt may favor sodium reabsorption and also may prevent massive potassium secretion along the CD in NS.

## Methods

### Animal experiments

The Institutional Ethical Committee of Animal Care of the University of Geneva and Cantonal authorities approved all animal experiments described in this work. The work described was carried out in accordance with the European-Union directive 2010/63EU. POD-ATTAC transgenic mice were a kind gift from Dr. P. Scherer (UT Southwestern Medical Center, Dallas, TX). They were bred and housed in the animal facility of the Faculty of Medicine of Geneva. Genotyping of transgenic mice was performed by PCR analysis of ear biopsies using the following primers: forward, 5’-GAA AGT GCC CAA ACT TCA GAG CAT TAG G – 3’ and reverse, 5’-AAC TGA GAT GTC AGC TCA TAG ATG GGG G-3’. These mice express a FKBP/Caspase-8 fusion protein under the control of the podocin promoter. NS was induced by intraperitoneal injection of 0.5 μg/g body weight of the chemical AP20187 (Takara Bio Inc. Kusatsu, Japan), in 8-10 weeks-old male transgenic mice. This compound induced dimerization of the fusion protein and caspase 8 activation resulting in podocyte-specific and glomerular proteinuria ^12^. Male wild-type littermates were used as controls and also received the same dose of AP20187. On the 2nd, 3rd or 5th day after AP20187 injection, mice were anesthetized by 100 mg/kg ketamine and 5 mg/kg xylazine (Bayer Healthcare, Berlin, Germany) and their kidneys were harvested before sacrifice by lethal bleeding. One kidney was immediately fixed by immersion in 4% paraformaldehyde and the second kidney was immediately frozen in liquid nitrogen before extraction of RNA and protein.

To assess the effect of sodium diet, male C57B6 mice (Charles River, Saint Germain de l’Arbresle, France) were fed for 7 days with either low-Na^+^ diet [0.01% (wt/wt); LSD] or normal-Na^+^ diet [0.18% (wt/wt); NSD] (Provimi-Kliba, Kaiseraugst, Switzerland). Animals had free access to food and water. Water deprivation experiments were performed as previously described ^13^. Male C57B6 mice were divided into two groups, control animals with access to food and water *ad libitum* and animals water deprived for 24 h.

ENaC blockade experiments were performed by the administration of the collecting duct diuretic amiloride (ENaC blocker) during 5 days. Amiloride solution (150 mg/L) was given in the drinking water to one group of male POD-ATTAC mice as previously described ^14^. Amiloride treatment was initiated after AP20187 injection to prevent massive sodium retention. Mice had free access to water and food throughout the study, and their water and food intake were measured during the experiment.

As previously described ^15^, NS in rats was induced by a single intra-jugular injection of 150 mg/kg body weight of puromycine-aminonucleoside (PAN) (Sigma–Aldrich, Saint Louis, MI, USA). Control rats received a single injection of 1 ml /100 g body weight of isotonic NaCl. Animals were anesthetized by 86 mg/kg ketamine (Vibrac, France) and 13.2 mg/kg xylazine (Bayer Healthcare, Berlin, Germany) 6 days after injection (the point of maximum sodium retention and proteinuria in PAN rats) and their kidneys were harvested before sacrifice by lethal bleeding. Kidneys were either fixed by immersion in 4% paraformaldehyde for histological analysis or frozen in liquid nitrogen and stored at −80°C for subsequent RNA and protein extraction.

### Human study

Human renal tissues were selected from the Archive of the Nephropathology unit (Service of Clinical Pathology, University Hospital of Geneva). The work presented has been carried out in accordance with The Code of Ethics of the World Medical Association (The Declaration of Helsinki) under the ethical committee approval CEREH number 03-081. Each patient provided informed consent before enrollment. Formaldehyde-fixed human renal tissues were dehydrated and paraffin-embedded. Five control renal tissues were obtained from distant peritumoral tissue obtained after nephrectomy performed for neoplasia and 5 kidney biopsies were obtained from patients with focal and segmental glomerulosclerosis (FSGS).

### RNA extraction and real-time qPCR

RNA extraction and real-time qPCR were performed as previously described ^16^. The primers used are listed in Table 1.

**Table 1:**
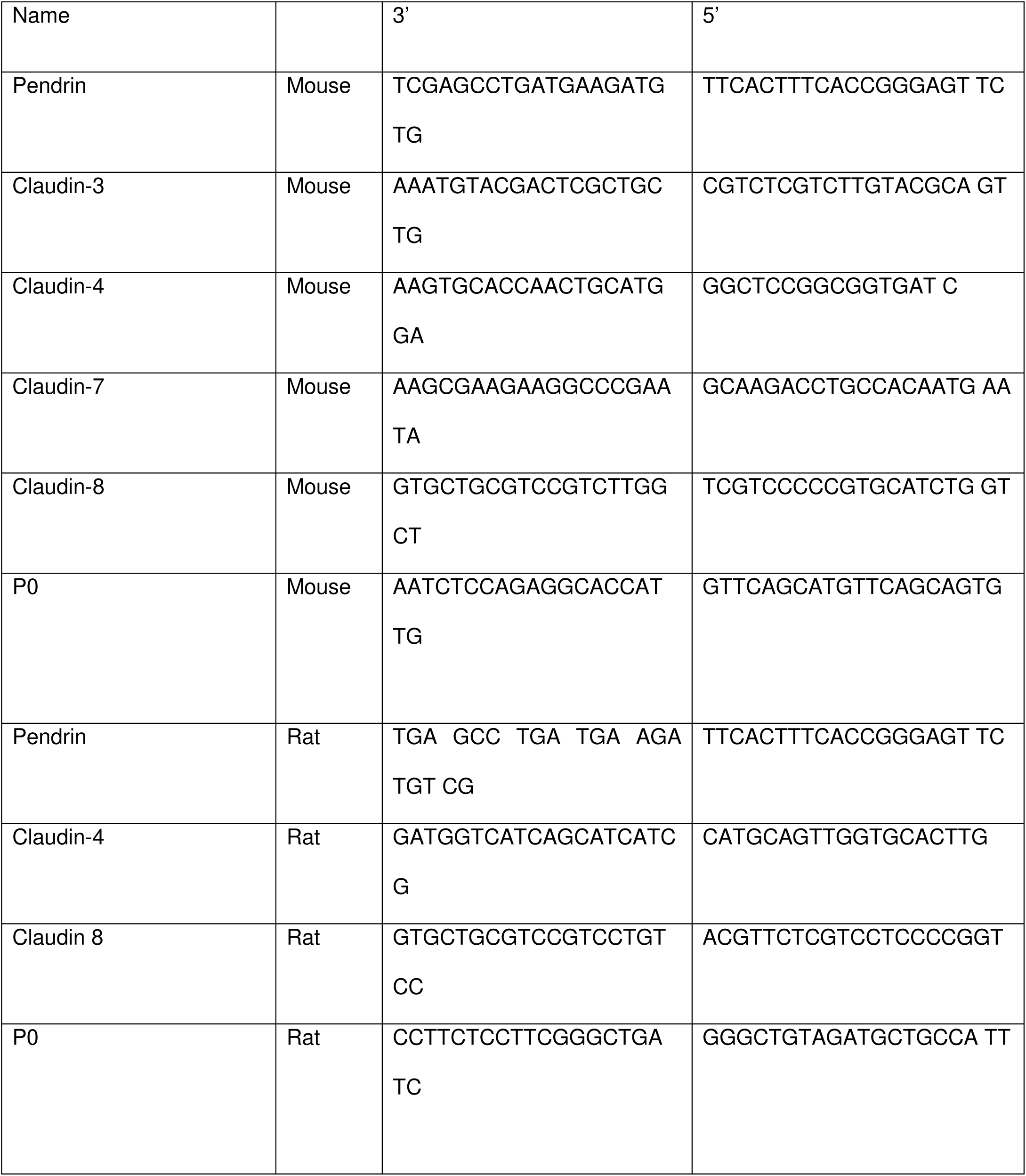
Primers for qRT-PCR.

### Western blotting

Kidney tissues or cultured cells were lysed as previously described ^17^. Proteins were quantified using a BCA protein assay kit (Pierce, Rockford, Il), subjected to SDS-PAGE and blotted onto polyvinylidene difluoride membranes (Immobilon-P; Millipore, Bedford, MA) using standard methods. The antibodies used are listed in Table 2. Detection of the antigen-antibody complexes was done by enhanced chemiluminescence using a PXi apparatus (Integrated Scientific Solutions, San Diego, CA). Protein signals were quantified with image J software. Results are expressed as the ratio of the densitometry of the band of interest to the loading control.

**Table 2:** Antibodies for Western blots and immunofluorescence.

| Name | Species | Dilution | Supplier | Reference |
| --- | --- | --- | --- | --- |
| γ-ENaC | Rabbit | 1/1000 | StressMarq<br>Biosciences | Ref SPC405 |
| Pendrin | Rabbit | 1/1000 | C Wagner | <sup>35</sup> |
| AQP2 | Rabbit | 1/1000 | RW Schrier | <sup>36</sup> |
| AQP2 | Mouse | 1/1000 | Santa Cruz<br>Biotechnology | Ref sc-515770 |
| H<sup>+</sup>ATPase | Rabbit | 1/5000 | Abcam | Ref ab121989 |
| Claudin-3 | Rabbit | 1/500 | Thermofischer | Ref 34-1700 |
| Claudin-3 | Rabbit | 1/500 | Abcam | Ref ab15102 |
| Claudin-4 | Mouse | 1/500 | Thermofischer | Ref 32-9400 |
| Claudin-4 | Rabbit | 1/1000 | Novus | Ref NBP2-41187 |
| Claudin-4 | Rabbit | 1/500 | Cusabio | Ref CSB-PA001662 |
| Claudin-7 | Rabbit | 1/500 | Thermofischer | Ref 34-9100 |
| Claudin-8 | Rabbit | 1/500 | Thermofischer | Ref 40-0700Z |
| NCC | Rabbit | 1/1000 | J Loffing | <sup>37</sup> |
| E-Cadherin | Mouse | 1/4000 | BD Biosciences | Ref 610181 |
| β-Actin | Mouse | 1/10000 | Sigma | Ref A5441 |

### Cell culture, constructs, viral particle production, and cell transduction

mCCD_cl1_ cells were grown on permeable filters (Transwell^®^, Corning Costar, Cambridge, MA) to confluence in a 1:1 mixture of Dulbecco’s modified Eagle’s medium and F12 medium, as previously described ^18^. Potential difference and transepithelial resistance were measured in the absence of any drug (except otherwise noted) using a Millicell-ERS Volt-Ohm meter (Millipore, Billerica, MA).

For the gene overexpression, cells were transduced with empty lentiviruses (empty), wild-type green fluorescence protein (GFP), or that encoding wild-type mouse claudin-4. For gene silencing, claudin-4, or scramble short hairpin RNA (shRNA) (Table 3) was inserted into the plasmid pLKO.1 (8453; Addgene). pSF-lenti or pLKO.1 together with the two helper plasmids psPAX2 (Addgene plasmid 12260) and pMD2.G (Addgene plasmid 12259) were transiently co-transfected in packaging HEK293T cells using the Polyplus-transfection^®^ jetPRIME^®^ Kit according to the manufacturer’s instructions. Lentiviral particles were collected after 72 h and γ-ENaCTetOn-mCCD cells were transduced. Stable polyclonal cell lines were selected 72 h after transduction.

**Table 3.**
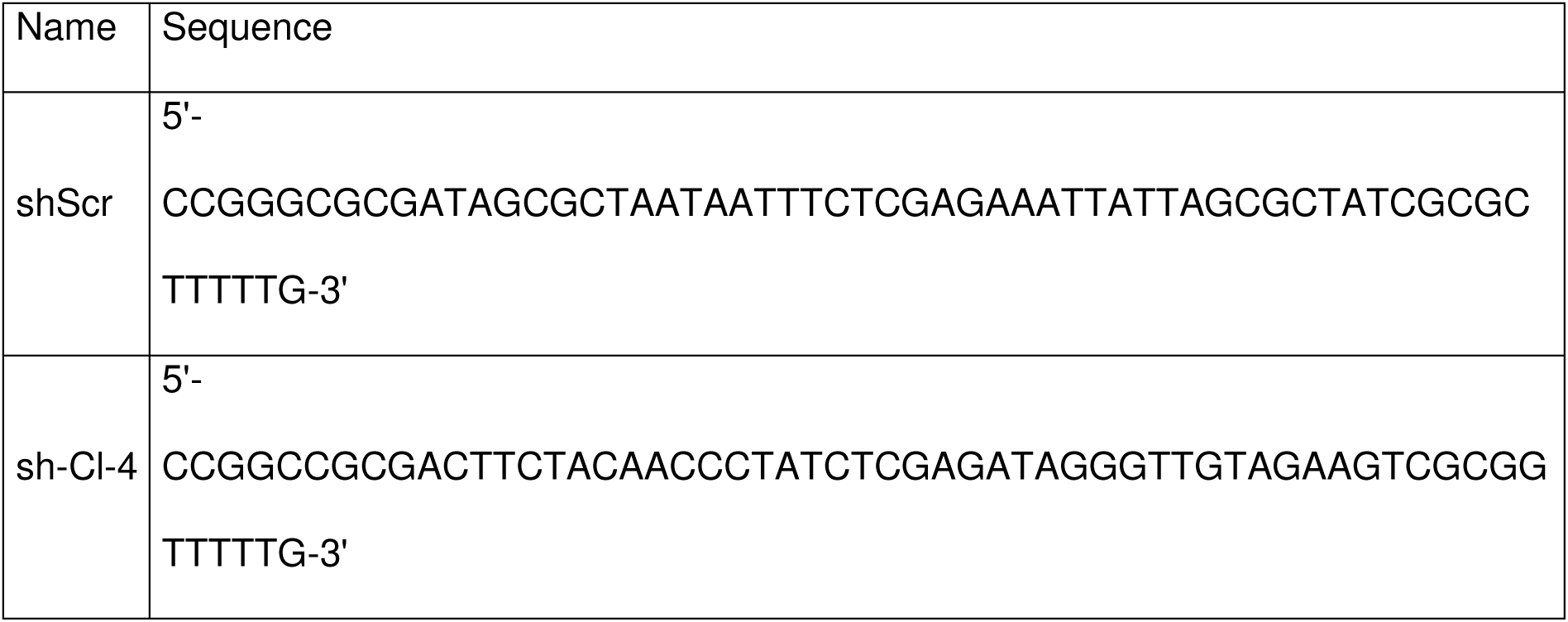
Sequences of scramble shRNA (shScr) and shRNA targeting claudin-4.

### Measurement of dilution potentials

Cultured mCCD_cl1_ cells were seeded onto 1.12 cm^2^ Snapwell polyester filters (Snapwell; Corning Costar, Cambridge, MA) (250,000 cells/cm^2^) and incubated in medium for 7 days. The Snapwell rings were then detached and mounted in Ussing chambers (model P2300; Physiologic Instruments, San Diego, CA, USA). The Ussing chambers were connected to a VCC MC6 multichannel voltage/current clamp (Physiologic Instruments) *via* silver/silver Cl^-^ electrodes and 3 M KCl agar bridges. The apical and basolateral hemi-chambers were separately filled with buffer A (120 mM NaCl, 10 mM NaHCO_3_, 5 mM KCl, 1.2 mM CaCl_2_, 1 mM MgCl_2_, and 10 mM Hepes, pH 7.4). The fluid volume on each side was 5 ml. Transepithelial potentials between the apical and basal hemi-chambers were recorded using a Quick Data Acquisition DI100 USB board (Physiologic Instruments) with the transepithelial current clamped at 0 mA during the whole experiment. Cells were equilibrated for 1 hour in buffer A. Dilution potentials were measured by replacing half the buffer in the basal side with buffer B (240 mM mannitol, 10 mM NaHCO_3_, 5 mM KCl, 1.2 mM CaCl_2_, 1 mM MgCl_2_, and 10 mM Hepes, pH 7.4) to measure the dilution potential of NaCl. Mannitol was used to maintain osmolarity and pH was adjusted to 7.4 with HCl. To ensure ion flux was occurring *via* a paracellular pathway, we added 100 µM amiloride (Sigma) and 100 μM 4,4ʹ- diisothiocyanatostilbene-2,2ʹ-disulfonic acid [DIDS] (Sigma-Aldrich) to the apical compartment for 30 minutes before performing dilution potential measurements. During all experiment, buffers were maintained at 37°C and bubbled constantly with a mixture of 95% oxygen and 5% carbon dioxide. Peak values of dilution potentials were measured and used to calculate the permeability ratios. The relative and absolute ion permeabilities were calculated as previously described ^19^. Briefly, the permeability ratio, *P*_Na_/*P*_Cl_ was calculated by using the Goldman–Hodgkin–Katz equation and absolute permeabilities for Na^+^ and Cl^−^ were calculated by using the Kimizuka-Koketsu equation from the calculated *P*_Na_/*P*_Cl_ and the transepithelial resistance measured during the same experiment.

### Immunofluorescence

After dehydration and paraffin-embedding, kidneys sections of 5 µm thickness for animal section and 3 µm thickness for human sections were used for analysis. Antigen retrieval was done with citrate buffer 10 mM pH6. Permeabilization with Triton 0.2% in PBS followed by blocking of non-specific binding site with bovine serum albumin 2% were applied for 30min. The tissue sections were then incubated 1h at 37°C with primary claudin-4 antibody (diluted 1:50 for animal sections, 1:10 for human sections), AQP2 antibody (diluted 1:400), H^+^-ATPase or pendrin antibodies (diluted 1:100, in PBS-NGS 10%) followed by a 30 min-incubation with a secondary Alexa Fluor 488-conjugated goat anti-rabbit (cat. no. A-11017; Invitrogen) or Cyanin3-conjugated goat anti-mouse (cat. no. M30010; Invitrogen) diluted between 1:200 to 1:800 in PBS-NGS 10% at 37°. Samples were mounted on microscope slides using Vectashield mounting medium (Maravai Life Science, San Diego, CA, USA) with DAPI for nuclear counterstaining. Fluorescence images were acquired using a Zeiss Axio Imager M2 (Carl Zeiss, Oberkochen, Germany). Negative controls were performed in the absence of primary antibody (not shown).

For cryosections, after Milestone Cryoembedding Compound (MCC) kidney tissue confection, tissue sections of 5 µm thickness were used for analysis. Fixation was done in methanol at − 20°C for 5 min. Subsequent steps were identical to those used for paraffin-embedded sections, as described above.

Cultured mCCD_cl1_ grown on polycarbonate filters were fixed with ice-cold methanol for 5 min at −20°C and then washed with PBS during 30 min. Blocking of nonspecific binding sites was done with 2% bovine serum albumin in PBS (PBS-BSA). Cells were then incubated overnight at 4°C with antibodies against claudin-4 diluted 1:500 in 0.2% BSA followed by a 1 h incubation with Alexa Fluor 488-conjugated goat anti-rabbit (cat. no. A-11017; Invitrogen) diluted 1:500 and were finally mounted on microscope slides using Vectashield mounting medium (Maravai Life Science, San Diego, CA, USA) with DAPI for nuclear counterstaining. Fluorescence images were acquired using a LSM 700 confocal laser-scanning microscope (Carl Zeiss, Oberkochen, Germany) using 488-nm ray lasers. The distance between the Z-slices was 0.25 µm. From 5 to10 Z-stack images were processed per sample using the ZEISS ZEN Imaging Software. ZEN 2.3 (Carl Zeiss, Oberkochen, Germany).

### Statistics

Results are presented as the mean ± SD from n independent experiments. Prism version 7.02 (GraphPad Software, San Diego, CA) was used for statistical analysis. The normal distribution of the population from which sample data was extracted was determined by a Shapiro-Wilk test. For data with normal distribution, statistical differences were assessed using a two-tailed unpaired Student t-test. For data with non-normal distribution, statistical differences were assessed using the non-parametric Mann-Whitney U-test. When more than two groups were compared, a one-way ANOVA with multipair wise comparison from Tukey was used for data with normal distribution whereas non-parametric Kruskal-Wallis test was used for data with non-normal distribution. A p-value < 0.05 was considered significant.

## Results

### Claudin-4 expression increases in parallel with cleaved γ-ENaC abundance in experimental nephrotic syndrome

As previously described, POD-ATTAC mice developed a typical NS defined by massive proteinuria, ascites, and hypoproteinemia ^16^. We have demonstrated massive sodium retention in this model and shown that the site of tubular sodium reabsorption shifted from the distal convoluted tubule *via* the sodium-chloride cotransporter NCC at day 2 after induction of the nephrotic syndrome to the collecting system (connecting tubule and CD) *via* ENaC from day 3 and later ^16^. Enhanced ENaC-dependent sodium reabsorption was correlated to the increased abundance of the cleaved active form of γ-ENaC. In parallel with sodium retention, nephrotic POD-ATTAC mice massively retained chloride (Fig.1A-B) calling for analysis of chloride transport pathway in NS.

**Figure 1.**
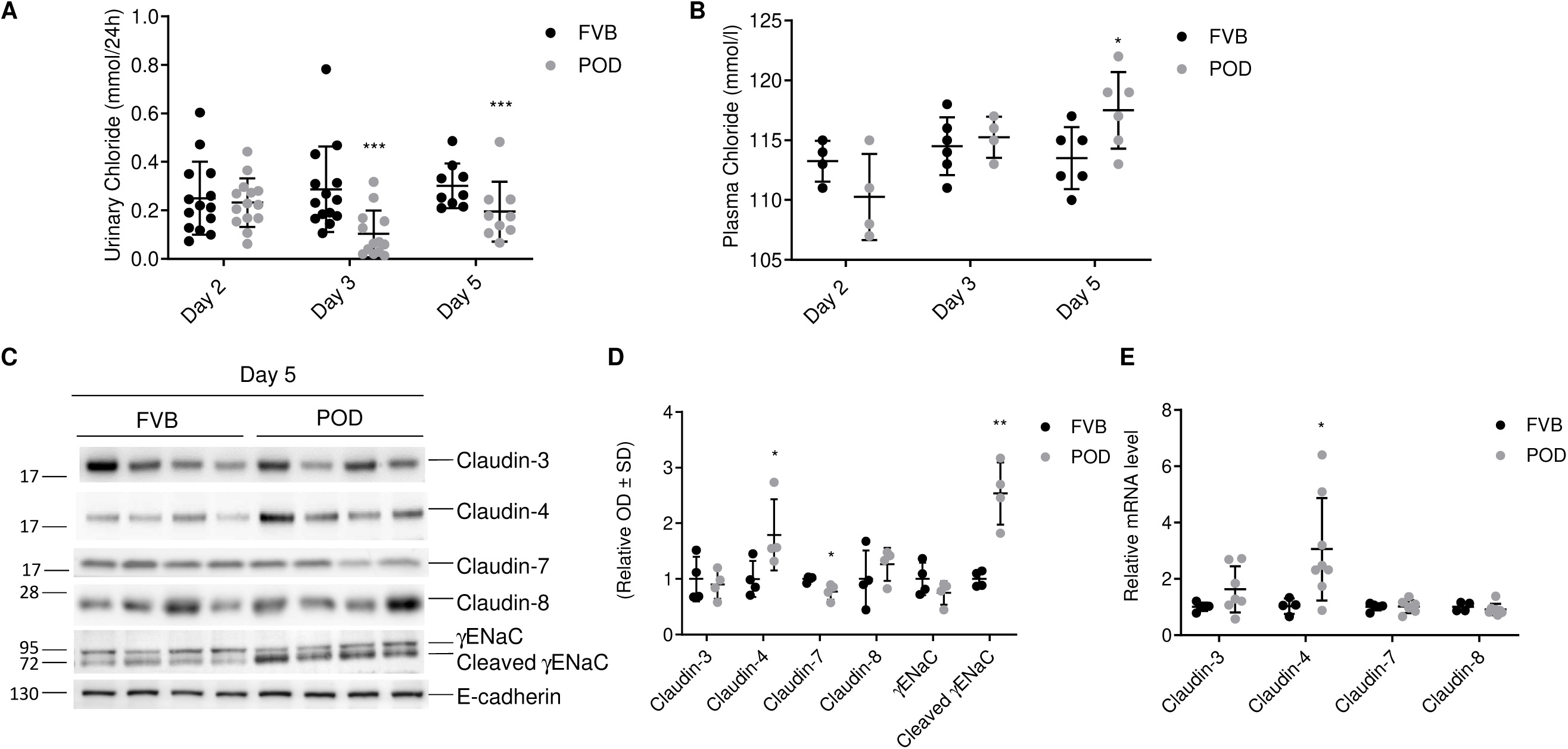
Time-course of γ-ENaC and claudins expression levels in nephrotic POD-ATTAC mice. Urinary chloride excretion (A) and plasma chloride (B) in control (FVB) POD-ATTAC (POD) mice, 2, 3 and 5 days after intraperitoneal injection of the dimerizer AP20187. Total protein and RNA from kidney cortices of control (FVB) and POD-ATTAC (POD) were extracted 5 days after induction of nephrotic syndrome by intraperitoneal injection of the dimerizer AP20187. (C) Representative Western blots showing the abundance of full-length and cleaved γ-ENaC, claudin-3, -4, -7, and -8. E-cadherin was used as a loading control. (D) Bar graphs showing the densitometric quantification of Western blots shown in (C). (E) RT-PCR analysis showing mRNA levels of claudin-3, -4, -7 and -8. Values are means ± SD from 4 to 8 mice in each group. Statistical differences between control and POD-ATTAC mice were assessed using a two-tailed unpaired Student t-test; *p < 0.05, **p < 0.01.

We recently described a potential coupling between transcellular and paracellular transport ^20^. Therefore, we analyzed by RT-PCR and Western blotting the time-course of the expression levels of the major claudin species expressed in the collecting system (namely 3, 4, 7 and 8). The abundance of claudin-3, -4, -7 and -8 mRNA and protein was unchanged at day 2 (Fig. S1A-C) and day 3 (Fig. S1D-F) after induction of the NS in POD-ATTAC mice. As previously shown, the amounts of cleaved form of γ-ENaC protein were significantly increased at day 3 (Fig. S1D-E). After 5 days of NS, however, a large increase in claudin-4 mRNA and protein abundance was observed (Fig. 1C-E). We also detected a slight but significant decrease in claudin-7 protein abundance without change in mRNA levels. The expression levels of claudin- 3 and -8 remained unchanged. At this time point, the amounts of the cleaved form of γ-ENaC remained elevated, confirming our past work ^16^.

We also studied the PAN rat as a reference model of NS. Six days after the induction of NS by injection of puromycin-aminonucleoside, rats displayed massive proteinuria and sodium retention ^21^. At this time point of NS, we identified an increase in claudin-4 and -8 protein abundance together with increased cleaved-fraction of γ-ENaC (Fig. 2A-B). This was accompanied by a non-significant rise of the mRNA levels of claudin-4 and a paradoxical decrease in claudin-8 mRNA levels (Fig. 2C).

**Figure 2.**
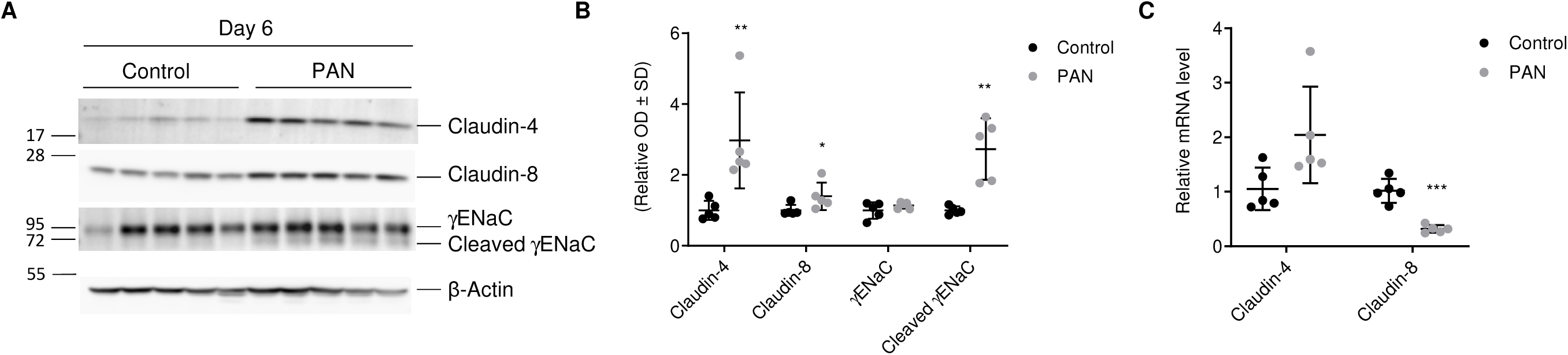
γ-ENaC and claudins expression levels in nephrotic PAN rats. Six days after intrajugular injection of vehicle (Control) or puromycin aminonucleoside (PAN) kidneys were harvested. (A) Representative Western blot experiment showing the abundance of full-length and cleaved γ-ENaC and of claudin-4 and -8. β-actin was used as a loading control. (B) Bar graph showing the densitometric quantification of Western blots shown in upper panel. (C) RT-PCR analysis showing mRNA levels of claudin-4 and -8. Values are means ± SD from 5 rats in each group. Statistical differences between control and PAN rats were assessed using a Mann-Whitney U test; *p < 0.05, **p < 0.01, ***p < 0.001.

These results obtained in two different rodent models show that claudin-4 expression levels are increased concomitantly with maximal sodium retention in NS.

### Claudin-4 expression in induced in collecting ducts and connecting tubules cells in experimental NS

We next analyzed the localization of claudin-4 along the kidney tubule by immunofluorescence imaging in control and nephrotic POD-ATTAC mice. We first assessed claudin-4 labeling in kidney cryosection. Panoramic views revealed a weak claudin-4 staining with polyclonal rabbit antibodies in control mice (Fig. S2A). Examination at large magnification revealed a typical reticulated pattern of claudin-4 staining indicating its localization along the tight-junctional complexes of CD (Fig. S2C). Double labeling with monoclonal mouse anti-AQP2 antibodies showed that claudin-4 positive tubules were cortical CD. The number of labelled tubules and the intensity of staining by anti-claudin-4 antibodies was dramatically increased in POD-ATTAC mice analyzed at day 5 after NS induction (Fig. S2B). AQP2 positive and some AQP2 negative tubules were positive for claudin-4 expression. These results were confirmed in PFA-fixed kidney sections, which much better preserves tissue architecture and cell shape integrity but disrupts tight-junctional complexes as shown by the loss of the typical reticulated claudin-4 staining replaced by a lateral membrane staining (Fig. 3). The very weak claudin-4 staining in control mice and the induction of claudin-4 expression in AQP2 positive and some AQP2 negative tubules in NS mice was also clearly observed in these preparations. Moreover, the better tissue preservation allowed us to show that most AQP2 positive and some AQP2 negative cells expressed claudin-4 in mouse CD. More precise analysis of the cellular localization of claudin-4 in mice was not possible because antibodies used to label claudin-4 and kidney tubule cell-specific markers were both obtained in rabbit, precluding their unambiguous separate detection by specie-specific secondary antibodies.

**Figure 3.**
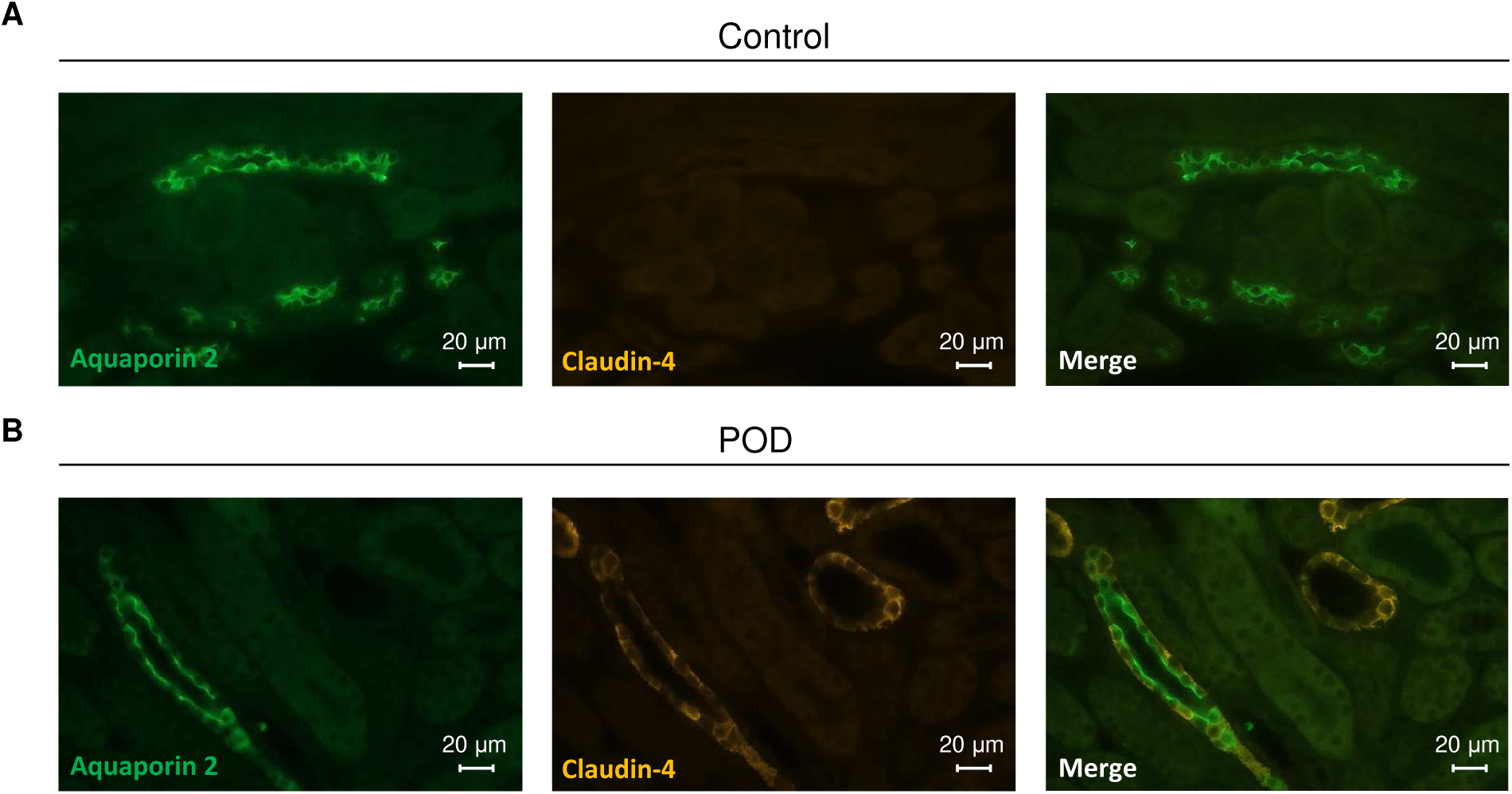
Claudin-4 localization in the collecting duct of control and POD-ATTAC mice. Mice kidneys were harvested and fixed in 4% paraformaldehyde 7 days after intraperitoneal injection of the dimerizer AP20187. Images show representative immunofluorescence double staining of claudin-4 with aquaporin-2 (AQP2) from 4 control (A) and POD-ATTAC mice (B).

To confirm our findings obtained in POD-ATTAC mice, we next assessed the localization of claudin-4 by immunofluorescence imaging in PFA-fixed kidney sections of control and PAN rats. The possible use of mouse anti-claudin-4 antibodies in rats combined with the well preserved structure of the kidney tissue fixed with PFA, allowed the detailed study of cellular localization of claudin-4 in control and PAN rats. In control rats, claudin-4 staining was weak both in panoramic views at low magnification (Fig. S3) and in focalized images at large magnification (Fig. 4A). In agreement with results obtained in POD-ATTAC mice, the number of cortical tubules stained by anti-claudin-4 antibodies increased in nephrotic PAN rats (Fig. S3). Strong claudin-4 staining was observed in pendrin-expressing B-IC (Fig.4B), and weaker claudin-4 labelling was present in AQP2 positive principal cells (Fig. 4A). However, claudin-4 staining was not observed in A-IC displaying apical H^+^-ATPase labelling in PAN rats (Fig. 4C). Claudin-4 staining was not observed in distal convoluted tubules identified by apical NCC staining (Fig. S4).

**Figure 4.**
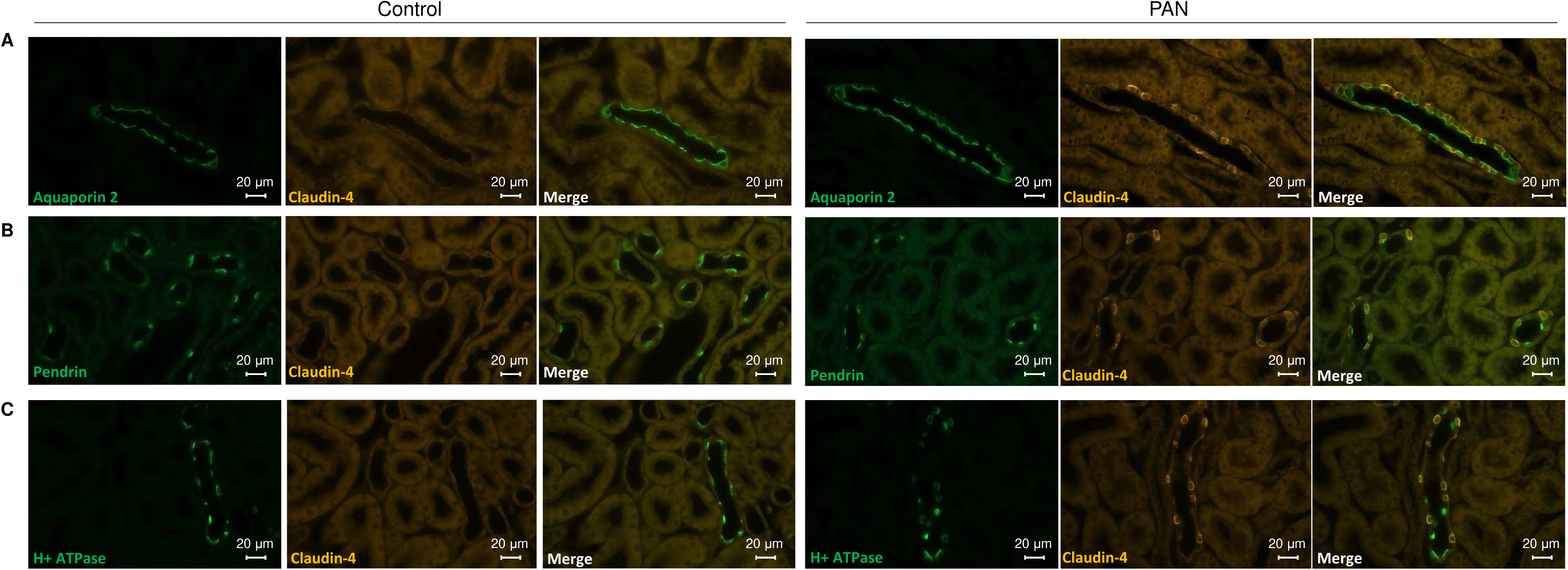
Claudin-4 localization in the collecting duct of control and PAN rats. Rat kidneys were harvested and fixed in 4% paraformaldehyde 6 days after intrajugular injection of vehicle (Control) or puromycin aminonucleoside (PAN). Images show representative immunofluorescence double staining of claudin-4 with aquaporin-2 (AQP2) (A), pendrin (B) and H^+^-ATPase (C), from 4 control and PAN rats.

These data indicate that claudin-4 expression is induced in CD and connecting tubules of nephrotic rodents, resulting in a large increase in number of claudin-4 expressing cells.

### Claudin-4 expression is induced in the collecting system of nephrotic patients with focal and segmental glomerulosclerosis (FSGS)

In order to confirm the pathophysiological relevance of our results obtained in mouse and rat models, the localization of claudin-4 was assessed by immunofluorescence in both normal and nephrotic human kidneys. In control human kidneys, claudin-4 expression levels are likely low, since claudin-4 immunostaining was only weakly observed in lateral membranes of collecting ducts identified by apical labeling with anti AQP2 antibodies (Fig. 5A). In contrast, kidney biopsies from nephrotic patients with a histological diagnosis of FSGS, displayed marked claudin-4 staining in both AQP2 positive principal cells and AQP2 negative IC (Fig. 5B). These results indicate that the claudin-4 induction observed in experimental rodent models of NS is also observed in human NS, suggesting its pathophysiological relevance.

**Figure 5.**
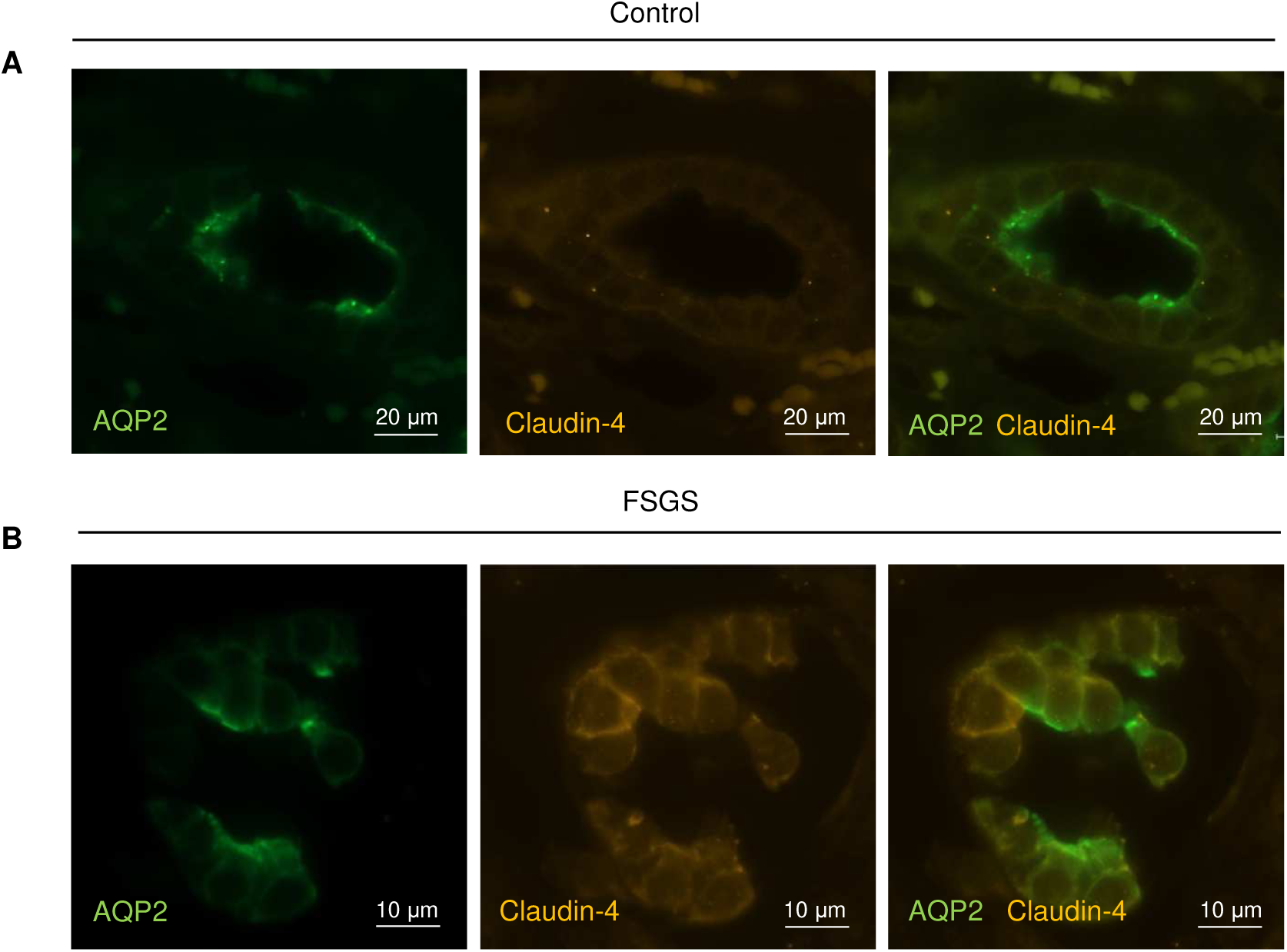
Claudin-4 expression is increased in collecting ducts of patients with nephrotic syndrome. Control (distant healthy tissue surrounding a kidney tumor) and nephrotic (kidney biopsies of patients with Focal and Segmental Glomerulosclerosis) human kidney pieces were fixed in formaldehyde before processing for indirect immunofluorescence analysis. Images show representative immunofluorescence double staining of claudin-4 with aquaporin-2 (AQP2) from 5 control (A) and nephrotic (B) humans.

### Increased claudin-4 abundance is associated with decreased pendrin expression levels in NS

Because claudin-4 may be involved in paracellular chloride transport along the collecting system in NS ^22^, we assessed the abundance of pendrin, the major chloride transporter in this part of the kidney tubule ^23^. Pendrin is an anion exchanger, which mediates the transcellular reabsorption of chloride and the secretion of bicarbonate by the B-IC of the CD ^24, 25^. Experiments performed after the induction of NS in POD-ATTAC mice did not reveal any change in pendrin protein abundance or RNA levels at day 2 (Fig. S5A-C). At day 3, protein levels were also unchanged (Fig. S5D-E), but a significant decrease in pendrin mRNA levels was measured in NS (Fig. S5F). At day 5, pendrin protein (Fig. 6A-B), but not mRNA levels were reduced in NS (Fig. S5G). In PAN rats, both pendrin protein and mRNA levels were significantly reduced 6 days after the induction of NS (Fig. 6C-D and Fig. S5H).

**Figure 6.**
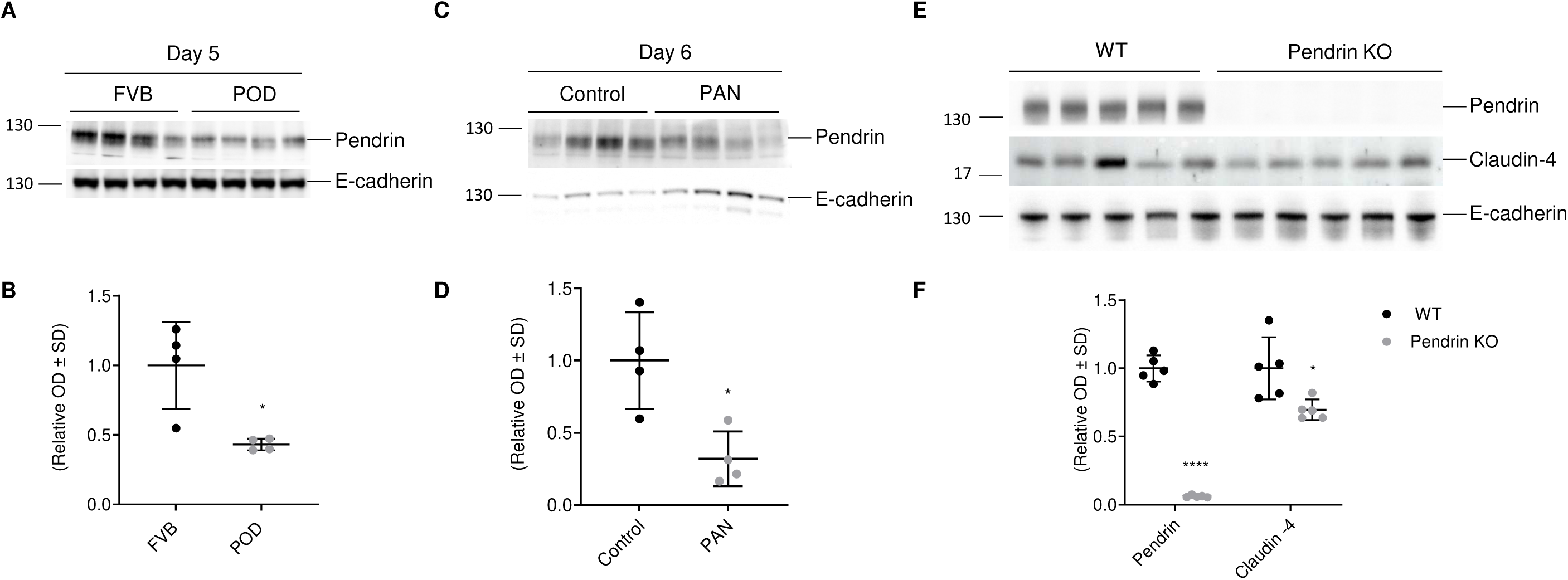
Pendrin expression in nephrotic POD-ATTAC mice and PAN rats. Total protein were extracted from kidney cortices of control (FVB) and nephrotic POD-ATTAC (POD) 5 days after intraperitoneal injection of the dimerizer AP20187 or 6 days after intraperitoneal injection of vehicle (Control) or puromycin aminonucleoside (PAN) in rats. A-B, Representative Western blot experiments showing the abundance of pendrin in FVB and POD-ATTAC mice. C-D Representative Western blot experiment showing the abundance of pendrin in control and PAN rats. E-F, Representative Western blot experiments showing the abundance of pendrin and claudin-4 in control (WT) and total pendrin knockout (pendrin KO) mice. E-cadherin was used as a loading control. Values are means ± SD from at least 4 animals in each group. Statistical differences between groups were assessed using a two-tailed unpaired Student t-test; *p < 0.05, ****p < 0.0001.

According the results described above, we tested the hypothesis that decreased pendrin expression is coupled to increased claudin-4 expression. We assessed claudin-4 abundance in global pendrin knockout mouse model ^26^ and showed that claudin-4 abundance is actually decreased in this setting (Fig. 6E-F). Therefore, pendrin and claudin-4 expression levels are not always inversely regulated and decreased pendrin expression does not explain increased claudin-4 expression in NS.

### Claudin-4 expression is stimulated by water deprivation in mice

POD-ATTAC mice display increased hematocrit and hemoglobin levels (Fig. S6), indicating an intravascular fluid volume depletion. We then decided to investigate the effect of vasopressin that is secreted in response to hypovolemia, on claudin-4 expression. We examined whether stimulation of endogenous vasopressin secretion by water restriction affects the expression of claudin-4. As expected, water deprivation increased AQP2 protein abundance, the vasopressin-regulated water channel (Fig. 7A-C). Claudin-4 protein abundance was increased in water-deprived mice as compared to control mice (Fig. 7A-C). This finding indicates that vasopressin may stimulate claudin-4 expression either directly *via* cAMP signaling or indirectly *via* stimulation of ENaC.

**Figure 7.**
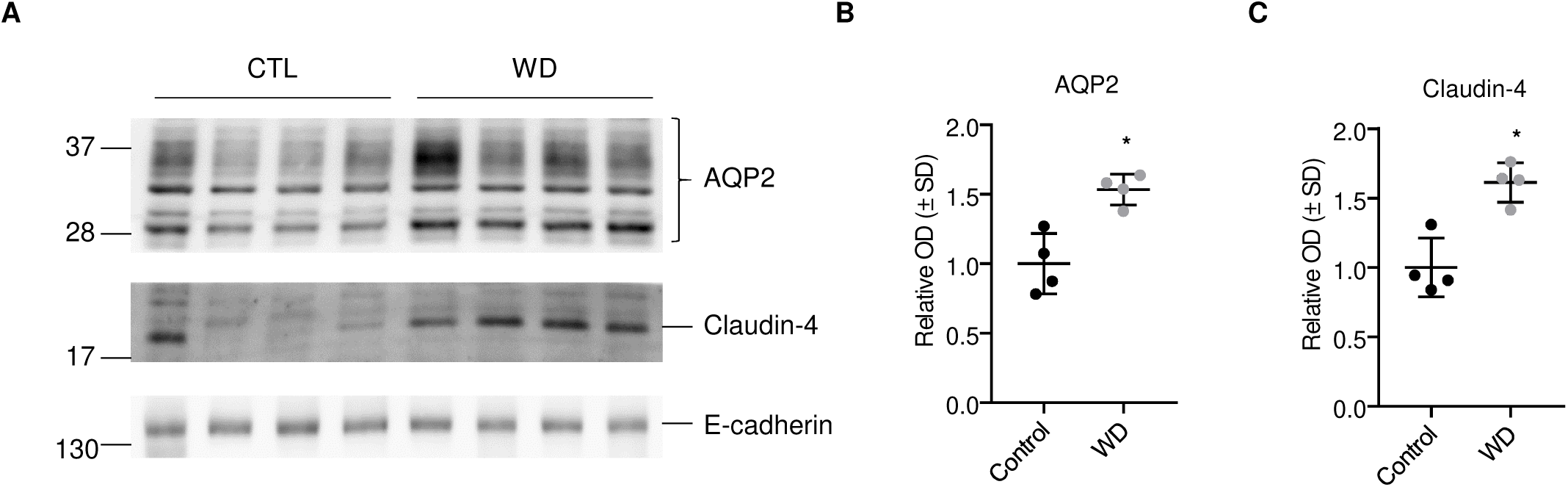
Claudin-4 expression is stimulated by water deprivation in mice. Effect of water deprivation on claudin-4 expression in normal mice. Control animals had access to food and water *ad libitum* and water-deprived mice (WD) were deprived of water for 24 h. A, Western blot analysis showing the effect of water deprivation on claudin-4 and AQP2 protein abundance on kidney cortex of 4 animals for each experimental group. B-C, Relative densitometric quantification of immunoblots shown in (A). E-cadherin was used as a loading control. Statistical differences between groups were assessed using a two-tailed unpaired Student t- test; *p < 0.05.

### Increased claudin-4 abundance is associated with increased ENaC activity and ascites formation in NS

In order to assess whether the observed change in claudin-4 abundance in POD-ATTAC and WD mice is mediated by enhanced ENaC activity, we administered amiloride (ENaC blocker) to POD-ATTAC mice for 5 days. *In vivo* administration of amiloride prevented ascites formation in nephrotic mice, as shown in Figure 8A-B by decreased body weight change and ascites quantity in amiloride-treated POD-ATTAC mice as compared to control mice. The cleaved active form of γ-ENaC protein abundance was increased in amiloride treated mice (Fig. 8C-D) suggesting that amiloride may increase endogenous vasopressin secretion and then stimulates ENaC expression. In contrast, claudin-4 induction observed in nephrotic mice was prevented by amiloride administration (Fig. 8C-D). These results indicate that claudin-4 induction in POD-ATTAC mice likely relies on increased ENaC activity.

**Figure 8.**
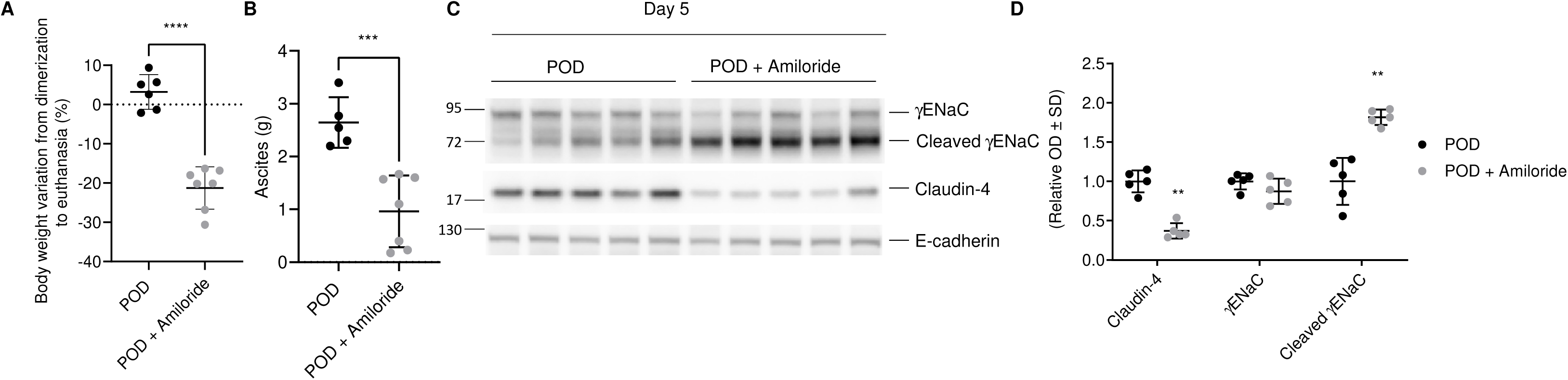
Effect of ENaC blockade in POD-ATTAC mice and claudin-4 expression. Amiloride (150 mg/L) was given in the drinking water during 5 days after induction of nephrotic syndrome by intraperitoneal injection of the dimerizer AP20187. (A-B) Effect of amiloride on body weight variation (A) and ascites quantity (B). (C) Representative Western blots showing the protein abundance of full-length and cleaved γ-ENaC and claudin-4. E-cadherin was used as a loading control. (D) Bar graphs showing the densitometric quantification of Western blots shown in (C). Values are means ± SD from 4 to 8 mice in each group. Statistical differences between control and POD-ATTAC mice were assessed using a two-tailed unpaired Student t-test; *p < 0.05, ***p < 0.001, ****p < 0.0001.

### Effect of claudin-4 overexpression on transepithelial resistance and paracellular transport in cultured CD principal cells

Our results in rodents and human suggest that claudin-4 may participate to the chloride retention of NS. To support this hypothesis, we assessed the physiological role of claudin-4. For this purpose, we constitutively overexpressed claudin-4 in mCCD_cl1_ cells. Real-time PCR (data not shown) and Western blot analysis (Fig. 9A-B) confirmed claudin-4 overexpression and the absence of altered abundance of the other major claudin species expressed along the CD, i.e. claudin-3, -7 and -8. Targeting of overexpressed claudin-4 to the tight junctions was confirmed by immunofluorescence imaging (Fig. 9C). Overexpression of claudin-4 resulted in a significant increase in transepithelial resistance under open-circuit condition (Fig. 9D). To determine the electrophysiological properties of the paracellular pathway in these cells, monolayers of control and claudin-4 overexpressing cells were mounted in Ussing chambers and the dilution potentials were monitored under current-clamp conditions. The magnitude of dilution potential (mV) was decreased in claudin-4 overexpressing cells compared with control cells after a 130 mM to 65 mM basolateral NaCl dilution with constant 130 mM apical NaCl (Fig. 9E-F). The paracellular permeability ratio for Na^+^ over Cl^−^ (P_Na_/P_Cl_) was decreased in claudin-4 overexpressing cells (Fig. 9G), indicating a decrease in the paracellular permeability to Na^+^ and/or an increase in the paracellular permeability to Cl^-^. The absolute permeabilities for Cl^-^ (P_Cl_) and Na^+^ (P_Na_) indicate that the decrease in P_Na_/P_Cl_ was mainly caused by an increase in P_Cl_ (Fig. 9H) since P_Na_ displayed no significant change (Fig. 9I). These results indicate that claudin-4 may participate to paracellular Cl^-^ reabsorption along the collecting duct.

**Figure 9.**
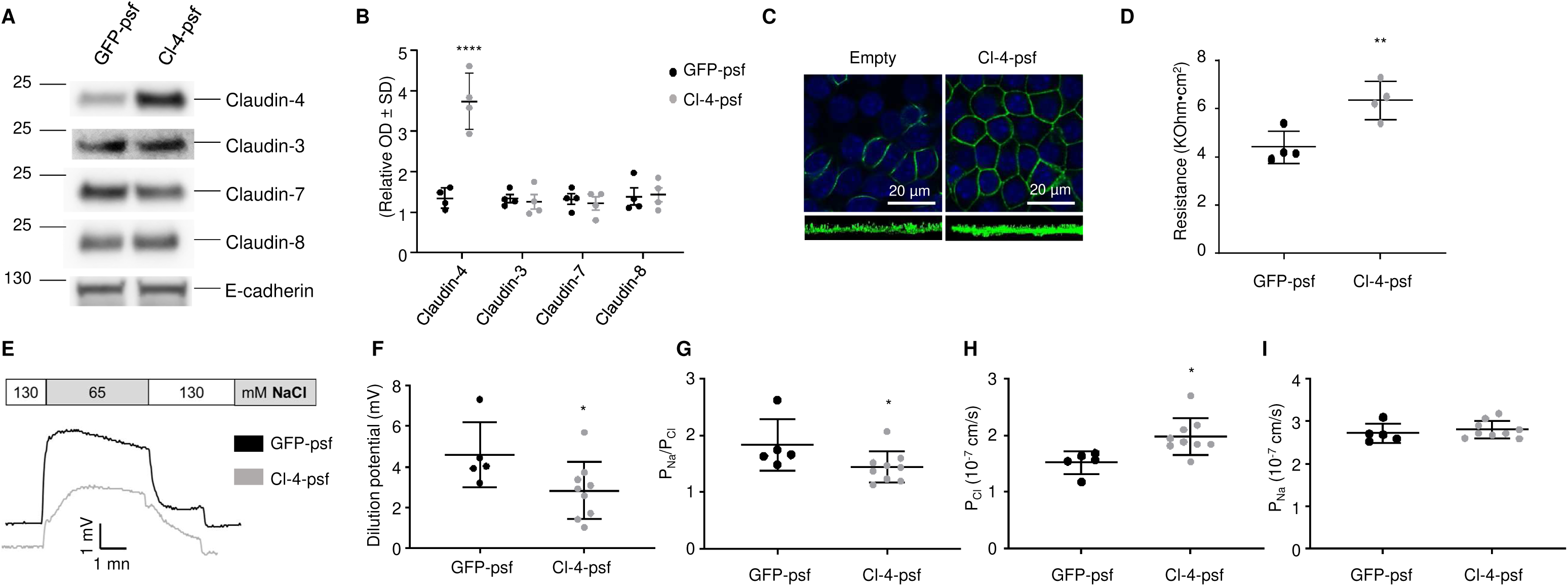
Effect of claudin-4 overexpression on transepithelial resistance and paracellular transport in cultured CD principal cells. mCCD_cl1_ cells were transduced with wild-type GFP lentiviruses (GFP-psf) or those encoding wild-type mouse claudin-4 (Cl4-psf). Transduced cells were grown to confluence on filters. (A) Representative immunoblots showing the effect of claudin-4 overexpression on claudin-3, claudin-7 and claudin-8 protein abundance. E-cadherin was used as a loading control. (B) Bar graphs showing the densitometric quantification of Western blots shown in (A). (C) Immunofluorescence staining of claudin-4 in confluent monolayers. mCCD_cl1_ cells transduced with empty lentiviruses (empty) were used as control. Lower part of each panel is an optical section obtained from Z-stack. (D) Transepithelial resistance (TER) measured under open-circuit condition. (E-F) Representative dilution potential recording in Ussing chambers under a zero-current clamp (E). The dilution potentials were determined from the change in transepithelial voltage upon switching the basolateral side from symmetrical bathing solutions (130 mM NaCl) to 65 mM basolateral NaCl (F). (G) Relative ion permeability ratios calculated from dilution potentials. (H-I) Absolute permeabilities of Cl^-^ and Na^+^ calculated from relative ion permeability ratios and the transepithelial resistance measured during the same experiment. Results are means ± SD from at least four independent experiments. Statistical analysis was performed by a Mann-Whitney U test; *p < 0.05, **p < 0.01, ****p < 0.0001.

### Effect of claudin-4 silencing on transepithelial resistance and paracellular transport in cultured CD principal cells

To confirm the function of claudin-4, we transduced cultured mCCD_cl1_ cells with shRNAs that specifically target claudin-4. ShRNA (Cl-4) that efficiently silenced claudin-4 was obtained as confirmed by Real-time PCR (data not shown), Western blot (Fig. 10A-B) and immunofluorescence (Fig. 10C). Claudin-4 did not alter protein abundance of claudin-3, -7 and -8 (Fig. 10A-B). Basal transepithelial resistance of cell monolayers, measured under open-circuit condition, was significantly decreased in cells transduced with sh-Cl-4 when compared with control cells (scramble) (Fig. 10D).

**Figure 10.**
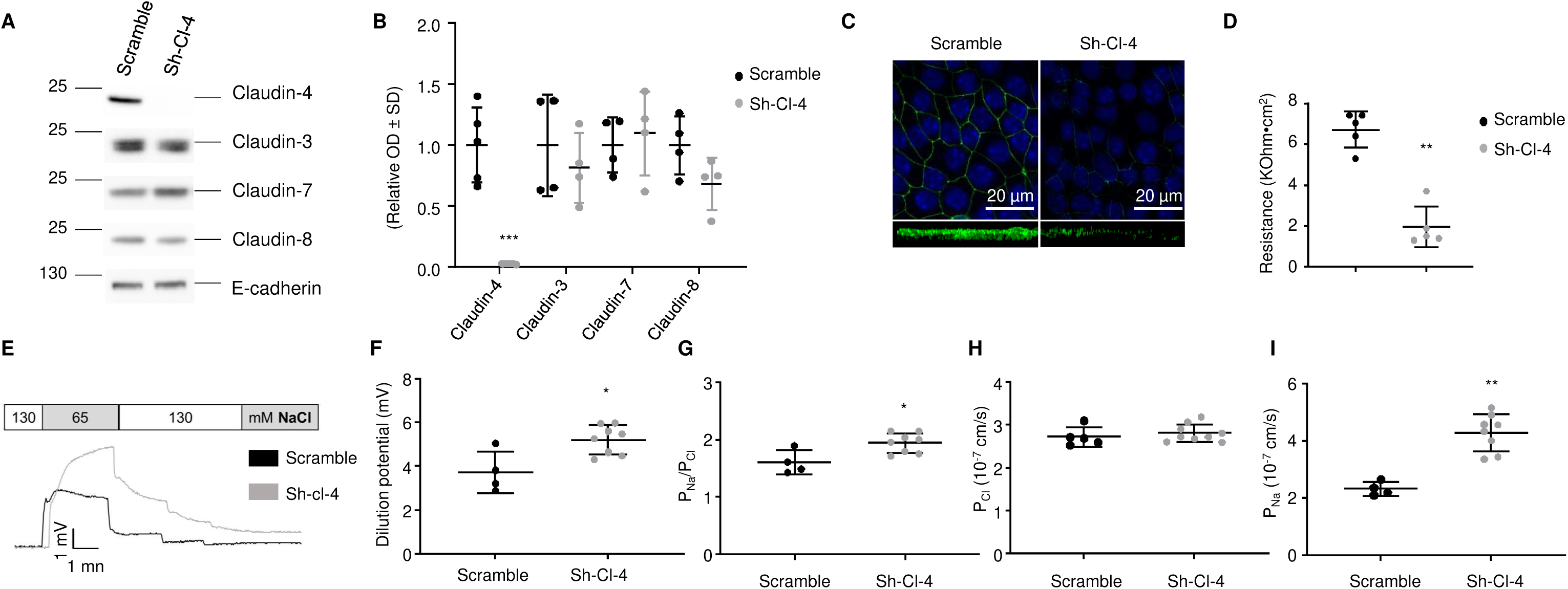
Effect of claudin-4 silencing on transepithelial resistance and paracellular transport in cultured CD principal cells. mCCD_cl1_ cells transduced with lentiviruses encoding either scramble shRNA (scramble) or shRNA targeting mouse claudin-4 (Sh-Cl-4) were grown to confluence on filters. (A) Representative immunoblots showing the effect of claudin-4 silencing on claudin-3, claudin-7 and claudin-8 protein abundance. E-cadherin was used as a loading control. (B) Bar graphs depicting relative densitometric quantification of immunoblots shown in (A). (C) Immunofluorescence staining of claudin-4 in confluent monolayers. Lower part of each panel is an optical section obtained from Z-stack. (D) Measured transepithelial resistance (TER) under open-circuit condition. (E-F) Representative dilution potential recording in Ussing chambers under a zero-current clamp (E). The dilution potentials were determined from the change in transepithelial voltage upon switching the basolateral side from symmetrical bathing solutions (130 mM NaCl) to 65 mM basolateral NaCl (F). (G) Relative ion permeability ratios calculated from dilution potentials. (H-I) Absolute pemreabilities of Cl^-^ and Na^+^ calculated from relative ion permeability ratios and the transepithelial resistance measured during the same experiment. Results are means ± SD from at least four independent experiments. Statistical analysis was performed by a Mann-Whitney U test; *p < 0.05, **p < 0.01, ***p < 0.001.

We then assessed the electrophysiological characteristics of mCCD_cl1_ cells displaying claudin-4 silencing. Figure 10E-F shows an increase in NaCl dilution potential in sh-Cl-4. The ion permeability ratio P_Na_/P_Cl_ was significantly increased in sh-Cl-4 transduced cells compared with control cells, indicating an increase in the paracellular conductance to Na^+^ and/or a decrease in the paracellular conductance to Cl^-^ (Fig. 10G). The absolute permeability for Na^+^ (P_Na_) was increased (Fig. 10I). In contrast, the silencing of claudin-4 did not alter the absolute permeability for Cl^-^ (P_Cl_) (Fig. 10H). These results suggest the possible role of claudin-4 as a barrier to Na^+^ along the collecting duct.

## Discussion

In our previous study using POD-ATTAC mouse, we demonstrated that sodium retention during NS initially takes place in proximal and distal tubules and quickly translates to the collecting system, where it relies on increased cleavage and apical localization of γ-ENaC ^16^. Here we showed that in NS, chloride reabsorption might be in part shifted from a transcellular energy consuming pendrin-dependent pathway to an energy free claudin-4-mediated paracellular pathway in the collecting system (connecting tubule and collecting duct).

Chloride transport along the collecting system is primarily mediated by pendrin, a Na^+^- independent Cl^-^ / HCO_3_ ^-^ exchanger ^24^, and is electrogenic when coupled to ENaC-mediated Na^+^ reabsorption by principal cells or electroneutral when coupled to NDCBE, a Na^+^-driven Cl^-^ / HCO_3_ ^-^ exchanger ^27^, located in the apical membrane of B-IC. Our results showed that pendrin protein abundance was decreased during the steady-state phase of sodium retention in both POD-ATTAC mice and PAN rats. This decrease in pendrin protein abundance was preceded by a decrease in its mRNA levels relying on either transcriptional down-regulation or reduced mRNA stability. These results suggest that at the time of massive sodium reabsorption in the CD ^16^, transcellular reabsorption of chloride is paradoxically down-regulated.

The down-regulation of the major transcellular chloride reabsorption pathway prompted us to analyze the changes in expression levels of the major claudin species expressed along the CD. Indeed, claudins determine the paracellular permeability to sodium and/or chloride through the tight-junction complexes. Results obtained in two rodent models of NS showed that mRNA and protein levels of claudin-4 were strongly increased. This up-regulation of claudin-4 was specific because the abundance of claudin-3 ^28^, claudin-7 ^29^ and claudin-8 ^20^ which are strongly expressed along the CD, was not systematically altered in NS. The induction of claudin-4 expression was confirmed by immunofluorescence imaging in both rodent models and in human NS. The expression of claudin-4 was strongly induced in both collecting ducts and connecting tubules but restricted to principal and β-intercalated cells. These results indicate that claudin-4 expression is inducible at least under pathological conditions of extreme sodium retention and confirms the plasticity of the composition of the tight-junctional complex.

Our *in vitro* experiments on mCCD_cl1_ cells confirmed that claudin-4 displays a preferential chloride over sodium pemeability^11, 22^. We have shown that overexpression of claudin-4 resulted in an increase in the paracellular conductance to Cl^-^ and that the silencing of claudin-4 increases the paracellular Na^+^ permeability, emphasizing the functional importance of claudin-4 as a Cl^-^ pore and Na^+^ barrier. In contrast, loss of claudin-4 did not detectably alter the paracellular permeability to Cl^-^ in transduced mCCD cells. This effect could be explained by the residual expression of claudin-4 detected by immunofluorescence imaging and/or the participation of other claudin species to the paracellular permeability to Cl^-^.

Taken together, our results strongly suggest that a paracellular chloride reabsorption pathway *via* claudin-4 is induced during NS. Chloride reabsorption *via* the paracellular pathway is driven by the lumen negative transepithelial voltage generated by ENaC-mediated sodium reabsorption. In NS, the lumen negative transepithelial potential increases dramatically ^30^, thereby enhancing the driving force for paracellular anion reabsorption. In addition, the decreased transcellular chloride transport consecutive to reduced pendrin abundance should increase the luminal chloride concentration which also favors paracellular chloride reabsorption. This “chloride shunt” may partly dissipate the lumen-negative transepithelial potential and thereby further enhance sodium retention and also may limit potassium secretion, thereby preventing hypokaliemia in NS. Our findings mirror the renal sodium-chloride wasting observed in claudin-4 knockout mice ^11^, highlighting the potential pivotal role of claudin-4 in chloride reabsorption. In a near future, inhibiting the paracellular pathway may represent an additional tool to oppose the sodium retention of NS patients. Currently available claudin-4 binders could weaken the tight junction barrier in opposition to ENaC-dependent sodium-chloride retention ^31^.

We have previously demonstrated that claudin-8 expression levels increased in parallel with ENaC γ-subunit abundance ^20^ demonstrating a coupling mechanism between transcellular and paracellular sodium permeability pathways. Since claudin-8 acts principally as a barrier to sodium, such a coordinated regulation prevents the backflow of reabsorbed sodium from the interstitial space to the tubular lumen. However, the present work does not confirm the presence of another coupling mechanism with an inverse correlation between pendrin and claudin-4 abundance to control chloride reabsorption. Indeed, claudin-4 expression levels are decreased in pendrin knockout mice ^32^. Therefore, pendrin deficiency is not obligatorily compensated by increased claudin-4 abundance.

We have recently shown that aldosterone increases claudin-8 but does not alter claudin-4 expression levels in cultured collecting duct principal cells ^33^. This finding suggests that induction of claudin-4 expression observed in nephrotic syndrome does not rely on increased aldosterone levels.

In POD-ATTAC mice, NS most likely increases non-osmotic vasopressin secretion *via* plasma volume contraction as suggested by increased hematocrit. This is supported by the observation of increased AQP2 expression in POD-ATTAC mice ^16^. This increased vasopressin secretion does not directly explain the induction of claudin-4 in NS that rather relies on stimulated ENaC activity. Indeed, inhibition of sodium retention by collecting duct diuretic, dramatically decreased ascites formation and claudin-4 induction, suggestion that the massive sodium reabsorption *via* ENaC in nephrotic syndrome is the main modulator of the paracellular sodium and chloride transport *via* claudin-4 regulation.

We have previously shown that mathematical modelling of POD-ATTAC mouse NS predicted an increased sodium transport and oxygen consumption in CNT/CD at day 5 compared to day 2 or 3. Shifting part of chloride transport from the transcellular to the paracellular pathway is expected to attenuate the increase in ATP and oxygen consumption by the CD. In support of this interpretation, Pei and colleagues showed that in the proximal convoluted tubule, the high proportion of paracellular solute reabsorption relying on claudin-2 optimized the energy efficiency and strongly reduced oxygen consumption by the kidney cortex, as compared to a purely transcellular reabsorption process ^34^. Altogether, these data suggest that chloride reabsorption along the ASDN shifts from an energy-demanding secondary active transcellular pathway *via* pendrin to an energy-saving passive paracellular pathway through claudin-4 in NS.

In summary, our data indicate that claudin-4 most likely plays an important role in paracellular chloride reabsorption accompanying ENaC-dependent sodium retention in nephrotic syndrome. We propose that claudin-4 inhibition may represent a novel target to remediate nephrotic edema.

## Data sharing statement

All data are available on reasonable request.

## Abbreviations

ASDN: aldosterone-sensitive distal nephron
AQP2: aquaporin-2
IC: intercalated cells
B-IC: β-intercalated cells
α-IC: α-intercalated cells
CD: collecting duct
CCD: cortical collecting duct
ENaC: epithelial sodium channel
NCC: sodium-chloride cotransporter
NS: nephrotic syndrome
PAN: puromycin aminonucleoside.

## Acknowledgements

We thank Mr. Thomas Cagarelli, Service of Clinical Pathology, University Hospital of Geneva, for processing the formaldehyde-fixed human renal tissues slides.

## Authors Contributions

V.O., E.F., A.S. and S.R. designed the study. V.O., A.S., G.A., A.C., I.R., K.U., E.D., S.M., J.M.R., L.C., G.C., C.A.W., A.D., R.C., D.E. and S.R. carried out experiments, V.O., A.S.,G.A.,S.R. and E.F. analyzed the data, S.R.,V.O.,A.S. and G.A made the figures, VO, AS, GA, S.R. and E.F. drafted and revised the paper, all authors approved the final version of the manuscript.

---

**Figure S1.**
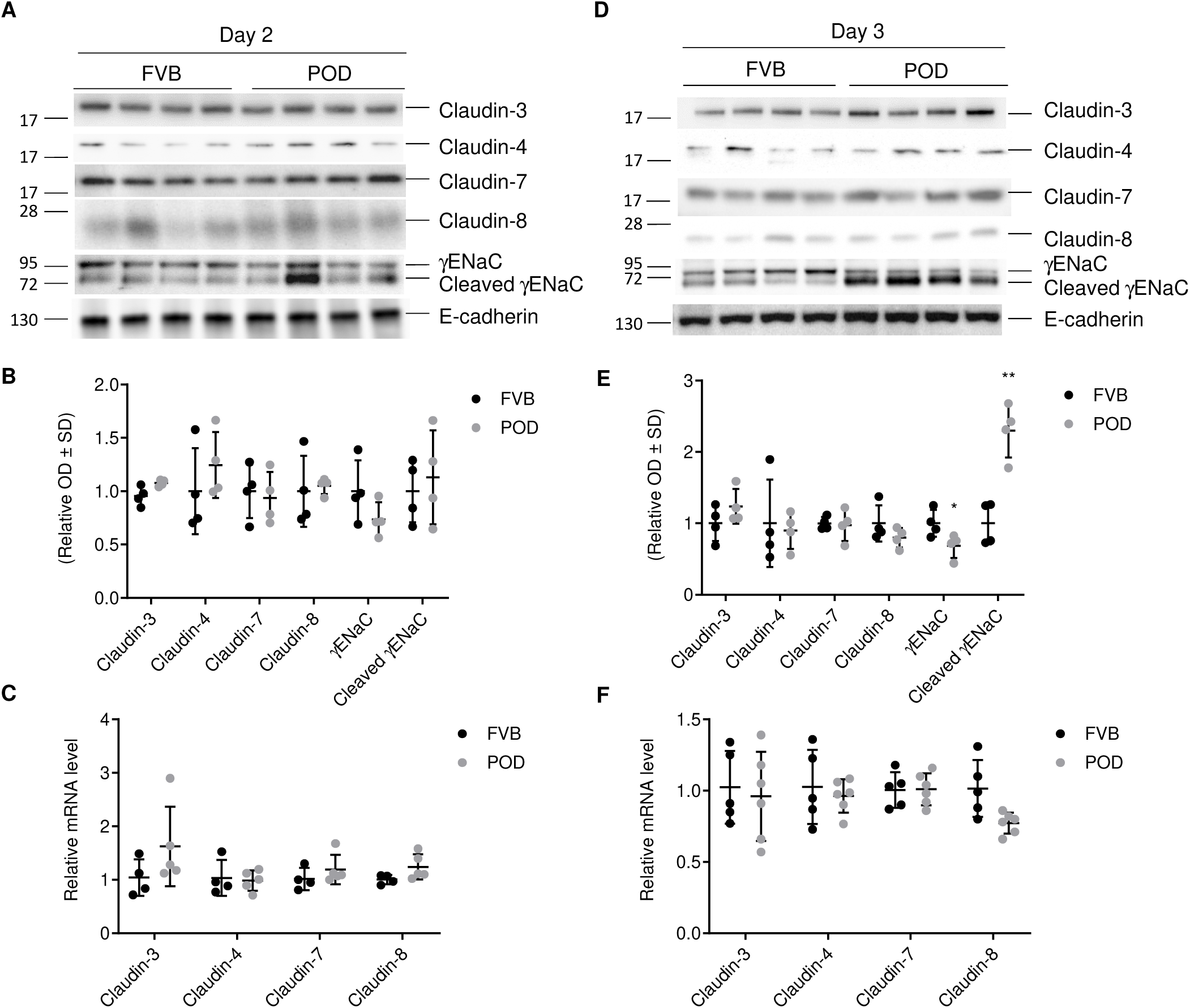
Time-course of γ-ENaC and claudins expression levels in nephrotic POD-ATTAC mice. Total protein and RNA from kidney cortices of control (FVB) and POD-ATTAC (POD) were extracted 2 or 3 days after induction of nephrotic syndrome by intraperitoneal injection of the dimerizer AP20187. (A) and (D) Representative Western blots showing the abundance of full-length and cleaved γ-ENaC, claudin-3, -4, -7, and -8. E-cadherin was used as a loading control. (B) and (E) Bar graphs showing the densitometric quantification of Western blots shown in upper panel. (C) and (F) RT-PCR analysis showing mRNA levels of claudin-3, -4, -7 and -8. Values are means ± SD from 4 to 8 mice in each group. Statistical differences between control and POD-ATTAC mice were assessed using a two-tailed unpaired Student t-test; *p < 0.05, **p < 0.01.

**Figure S2.**
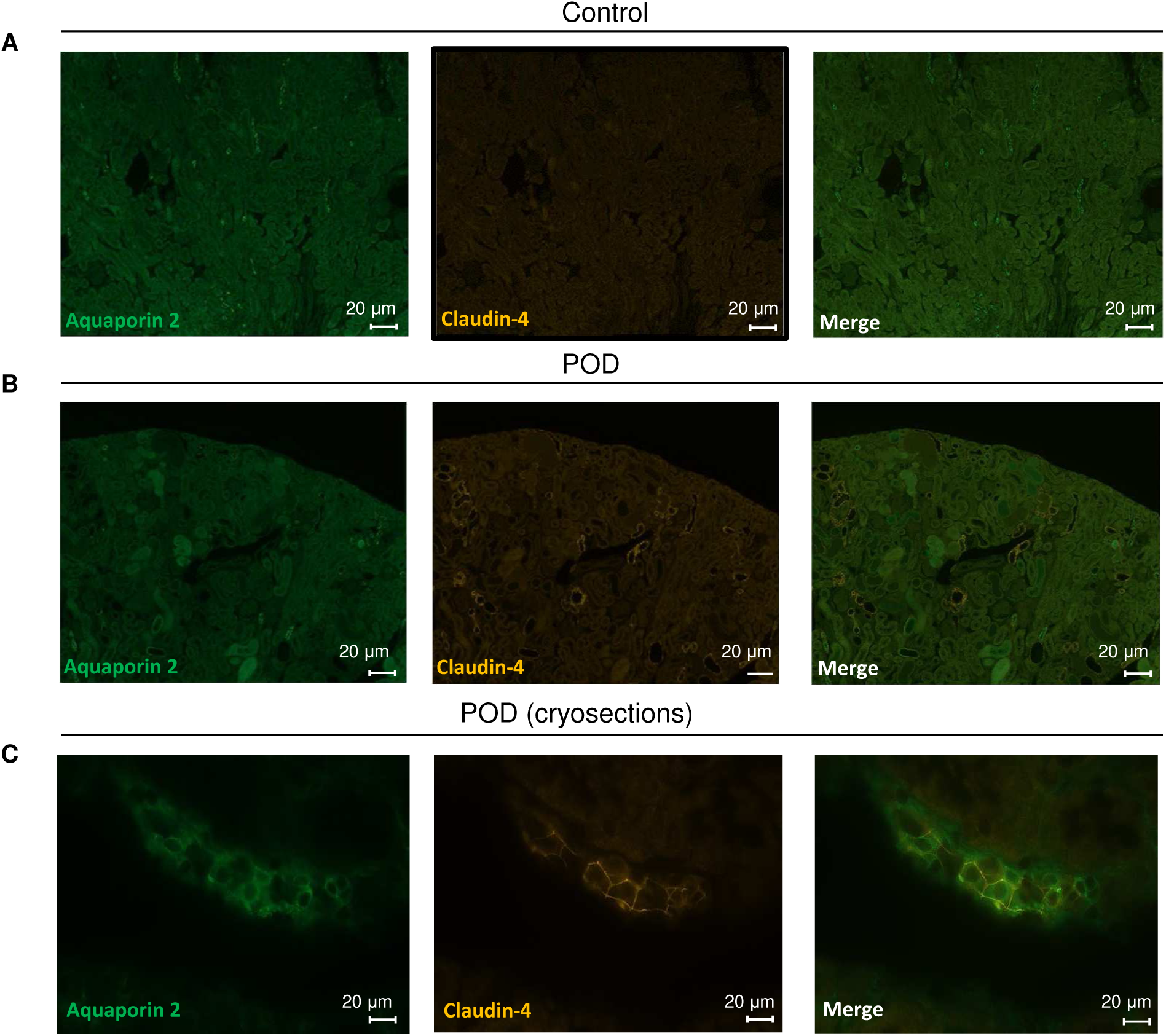
Claudin-4 localization in the collecting ducts of control and POD-ATTAC mice. Mice kidneys were harvested and fixed in 4% paraformaldehyde 7 days after intraperitoneal injection of the dimerizer AP20187. A-B representative panoramic views of double immunofluorescence staining of claudin-4 with aquaporin-2 in control (A) and POD-ATTAC mice (B). C, representative images of double immunofluorescence staining of claudin-4 with aquaporin-2 in cryosections from POD-ATTAC kidneys.

**Figure S3.**
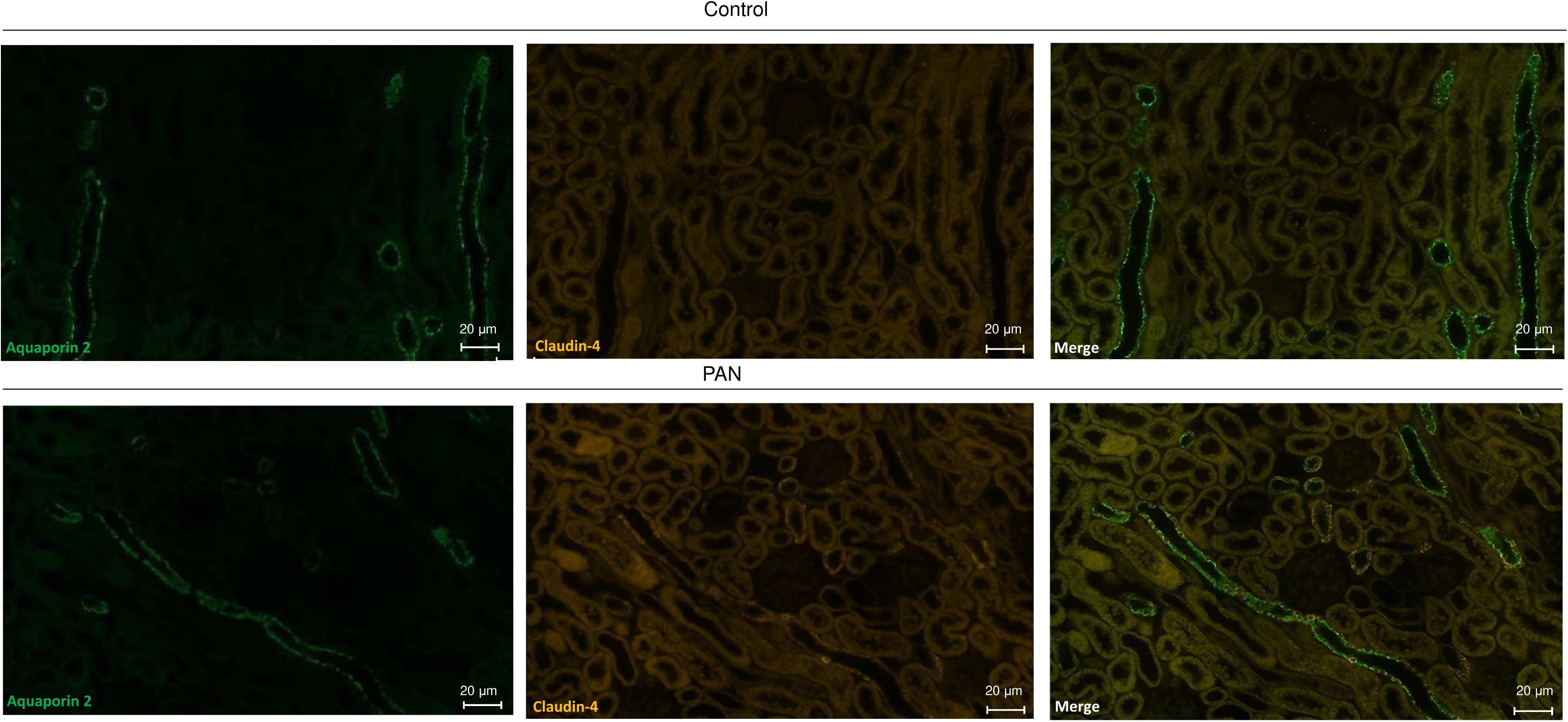
Claudin-4 localization in the collecting ducts of control and PAN rats. Rat kidneys were harvested and fixed in 4% paraformaldehyde 6 days after intrajugular injection of vehicle (Control) or puromycin aminonucleoside (PAN). Representative panoramic views of double immunofluorescence staining of claudin-4 with aquaporin-2 in control and PAN rats.

**Figure S4.**
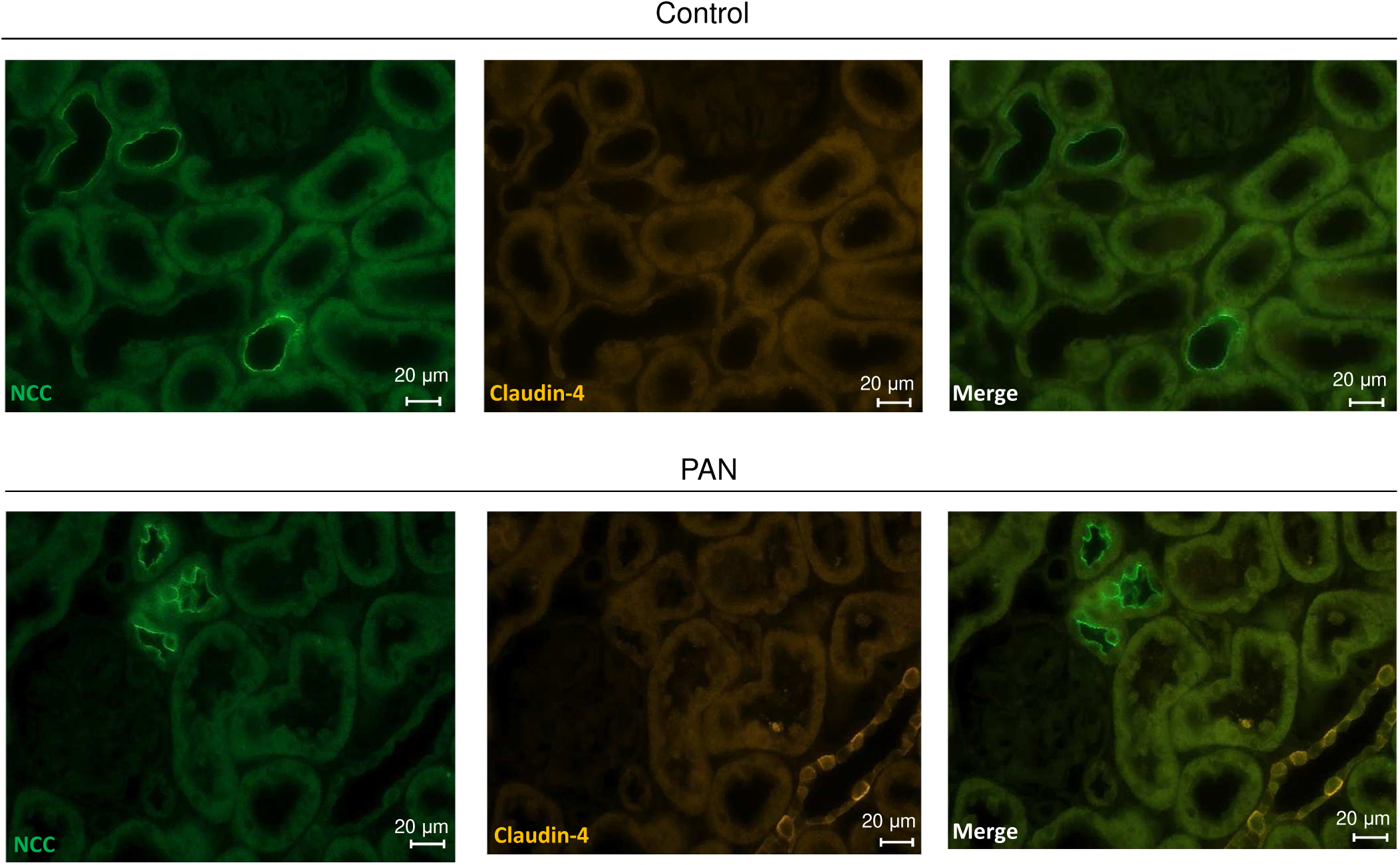
Claudin-4 localization in the collecting ducts of control and PAN rats. Rat kidneys were harvested and fixed in 4% paraformaldehyde 6 days after intrajugular injection of vehicle (Control) or puromycin aminonucleoside (PAN). Representative double immunofluorescence staining of claudin-4 with NCC in controls and PAN rats.

**Figure S5.**
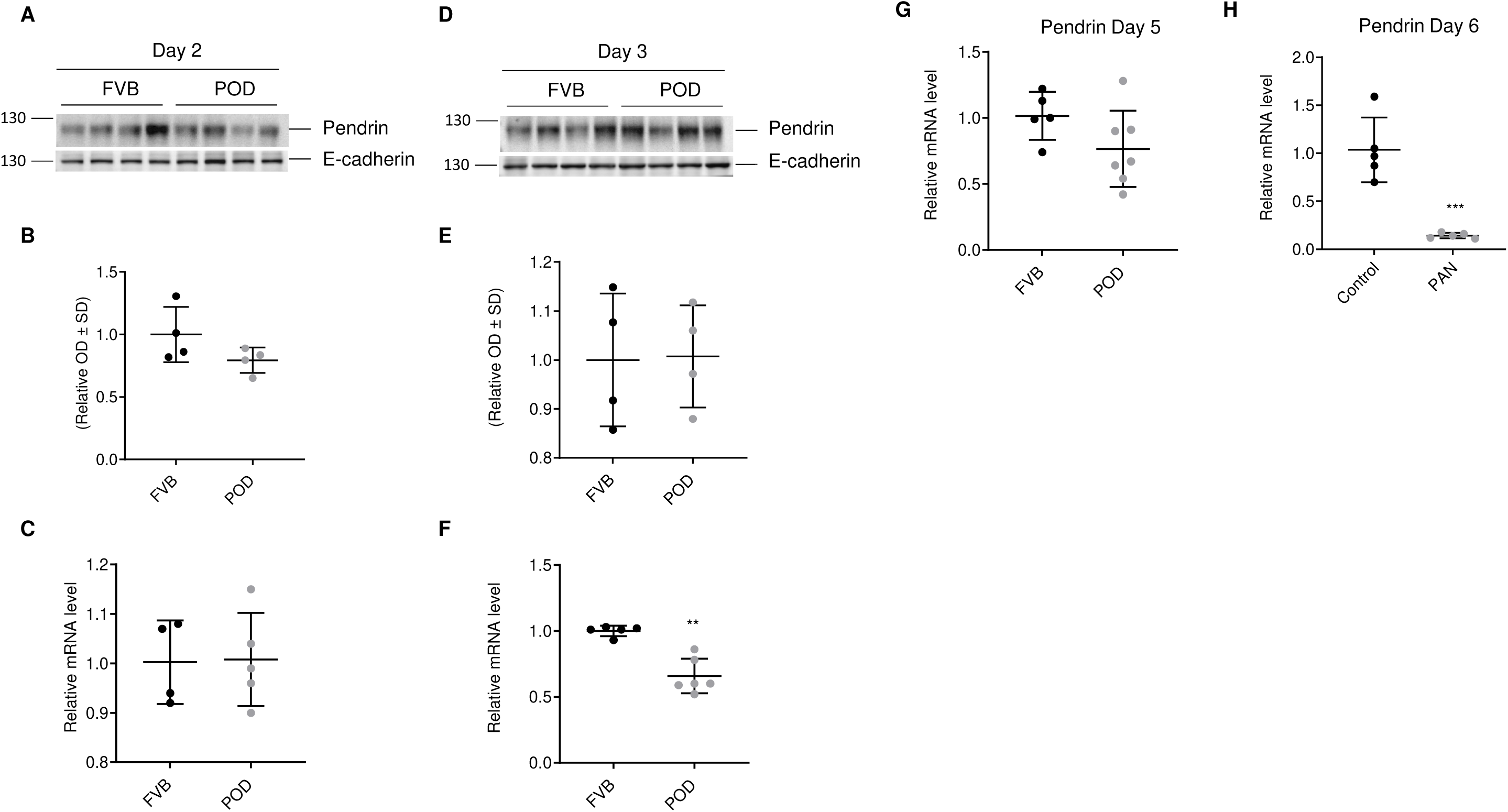
Pendrin protein and mRNA levels in POD-ATTAC mice and PAN rats. Total protein and RNA were extracted from kidney cortices of control (FVB) and nephrotic POD-ATTAC (POD) mice taken 2, 3 and 5 days after intraperitoneal injection of the dimerizer AP20187 or 6 days after intraperitoneal injection of vehicle (Control) or puromycin aminonucleoside (PAN) in rats. Representative Western blot experiments showing the abundance of pendrin in FVB and POD-ATTAC mice (A-B and D-E) and RT-PCR analysis of mRNA levels of pendrin in mice (C, F and G) and rats (H). Values are means ± SD. Statistical differences between groups were assessed using a two-tailed unpaired Student t-test; *p < 0.05, ***p < 0.001.

**Figure S6.**
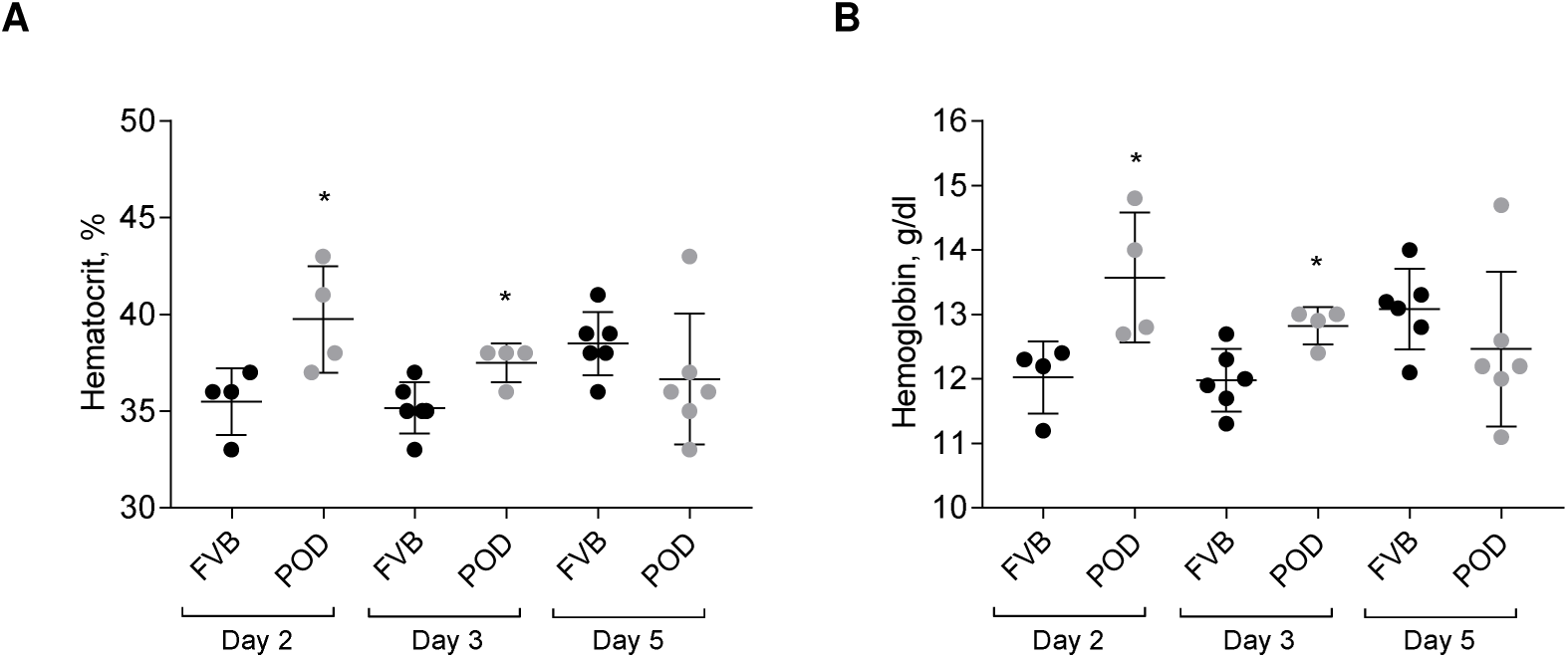
POD-ATTAC mice display hypovolemia. Hematocrit (A) and hemoglobin (B) in control (FVB) and POD-ATTAC (POD) mice, 2, 3 and 5 days after intraperitoneal injection of the dimerizer AP20187.

